# Analysis of *in vivo* infection dynamics using bioluminescent *Giardia duodenalis*

**DOI:** 10.64898/2026.03.02.709115

**Authors:** Rita Kosile, Bergeline Nguemwo-Tentokam, Peiran Shen, Nikki Farnham, Katherine McGowan, Scott Dawson, Steven M. Singer

## Abstract

*Giardia duodenalis* is an intestinal protozoan transmitted through contaminated food and water and is a major cause of diarrheal disease worldwide and has been linked to post-infectious sequelae and growth impairment in children. Quantifying parasite burden in vivo is essential for studying infection dynamics and host responses, yet commonly used methods (stool PCR, antigen detection, and terminal trophozoite counts from segments of the intestine) are limited by intermittent stool shedding, uneven parasite distribution, sampling error, and lack of longitudinal data. We developed a bioluminescent reporter system for the assemblage B strain GS to enable noninvasive tracking of infection in mice without antibiotic pretreatment. Using in vivo bioluminescent imaging (BLI), we observed peak signal in immunocompetent C57BL/6J and BALB/c mice at days 6–8 post-infection, followed by return to baseline by ~day 10, consistent with the self-limiting nature of giardiasis. In immunodeficient mice, radiance increased and persisted over time, demonstrating stable luciferase expression during chronic infection in vivo without continuous antibiotic selection. Whole-animal radiance strongly correlated with direct trophozoite counts from excised intestinal segments (r = 0.915, p < 0.001), validating BLI as a useful alternative for determining parasite burden. Together, these findings establish BLI as a robust platform for longitudinal studies of *Giardia* infection dynamics and for future applications that may examine host immunity, the microbiota, nutrient-dependent effects, and drug testing.

## INTRODUCTION

*Giardia duodenalis*, a protozoan parasite often spread through contaminated food or water, is one of the leading causes of diarrheal disease (Adam, 2001). The parasite exists in the environment in the form of a cyst. When ingested, it develops into trophozoites that colonize the upper small intestine by attaching to epithelial mucosa, where the trophozoites multiply (Dawson & House, 2010; Elmendorf et al., 2003). Trophozoites then encyst and are excreted back into the environment through the feces (Luján et al., 1997, 1998). Giardiasis is common in low- and middle-income countries, with an estimated 200 million cases reported annually. The global burden of giardiasis is uneven, affecting up to one-third of individuals in developing nations, compared to 3%–7% in more developed countries. The clinical presentation of giardiasis often includes diarrhea, abdominal cramping, and vomiting; however, a subset of infected individuals may remain asymptomatic (Kraft et al., 2017). In addition to acute symptoms, *Giardia* infection can result in post-infectious irritable bowel syndrome (Abedi et al., 2022; Hanevik et al., 2014), chronic fatigue syndrome (Hanevik et al., 2014, 2017), and growth stunting in children (Dougherty & Bartelt, 2022; Rogawski et al., 2018).

*Giardia* is classified into eight genetic assemblages (A–H) (Heyworth, 2016), with assemblages A (exemplified by the WB strain) and B (exemplified by the GS strain) being the most identified in humans (Feng & Xiao, 2011; Molina et al., 2011; Ramírez et al., 2015). The B assemblage of *Giardia* is a common genotype seen worldwide, but the dominant assemblage can vary by location and population studied (Feng & Xiao, 2011; Molina et al., 2011; Ramírez et al., 2015). These two assemblages are also the most widely utilized in animal and laboratory experiments. Other assemblages have been identified in livestock and companion animals (Durigan et al., 2014; Fantinatti et al., 2018; Volotão et al., 2007). Recently, several reports have shown human infections by assemblage E *Giardia* (Abdel-Moein & Saeed, 2016; Iwashita et al., 2021; Scalia et al., 2016; Zahedi et al., 2017). In vitro growth rate also varies between assemblages A and B, with assemblage B trophozoites exhibiting a slower in vitro growth rate. Assemblage B is also less efficient in excystation and encystation in vitro (Reiner et al., 2008).

Infection severity can vary across *Giardia* assemblages, but reports have been conflicting. While some studies have associated persistent diarrhea (Gelanew et al., 2007; Homan & Mank, 2001) or more chronic and sub-clinical infections (Ajjampur et al., 2009) with assemblage B parasites, other reports have shown that assemblage A parasites are associated with a higher incidence of diarrhea (Ajjampur et al., 2009; Haque et al., 2005; Sahagún et al., 2008). There have also been reports of no differences in symptomatology between assemblages A and B (Kohli et al., 2008; Zajaczkowski et al., 2021).

In the study of giardiasis, the use of in vivo models has been well established, and estimating the infection burden during specific experiments is essential. Several methods have been used to assess parasite burden. For example, DNA can be extracted from stool collected from infected animals or humans, and PCR then performed to quantify parasite DNA. Antigen ELISA can also be done to quantify infection burden in human stool samples (Duque-Beltrán et al., 2002). Additionally, mice can be euthanized and trophozoites counted directly by taking a small portion of the duodenum to determine the number of trophozoites per centimeter (Barash, Maloney, et al., 2017; Bartelt et al., 2013; Fink et al., 2020). These methods have certain limitations, including the potential for sampling errors. Stool shedding is known to be intermittent, potentially limiting the reliability of the PCR approach, and uneven distribution within the small intestine can lead to sampling errors when counting short segments of duodenum. Moreover, terminal parasite counts prevent the collection of longitudinal data.

The use of bioluminescent imaging (BLI) is becoming increasingly common, as it facilitates the visualization of a wide variety of *in vivo* cellular events without the need for euthanasia. This technique also enables repeated imaging and longitudinal monitoring of individual animals, thereby reducing variability and measurement errors. Barash, Nosala, et al., 2017 previously developed and applied BLI to investigate the *in vivo* dynamics of *Giardia* infection in live mice using a genetically modified strain of *Giardia* WB from assemblage A. This strain was engineered to express firefly luciferase (*FLuc*), offering a dynamic, noninvasive approach compared to traditional manual parasite counts. The gene encoding the *Fluc* was integrated into the WB genome under the control of either the cyst wall protein 1 (*CWP1*) promoter or the glutamate dehydrogenase (*GDH*) promoter. Both strains were used to infect antibiotic-treated C57BL/6J mice, and live imaging confirmed the bioluminescent activity of the parasites. The study showed that infecting trophozoites tend to cluster in specific areas of the small intestine in vivo. The intensity of these clusters also correlated with infection levels measured by stool PCR, though not with direct parasite counts. An important limitation of this study is that WB parasites require gnotobiotic mice or mice treated with high-doses of antibiotics to colonize the host successfully. Furthermore, while parasite burden in dissected segments of the small intestine was shown to be similar using both BLI and qPCR, no correlation was made between bioluminescent signals in whole animals and standard methods for quantifying infection burden.

We have now generated a genetically modified, bioluminescent strain of the assemblage B isolate GS. Unlike WB, GS can infect adult mice with an intact microbiome and has been used extensively to characterize immune responses, pathogenesis, and infection dynamics in this system (Bartelt et al., 2013; Byrd et al., 1994; Solaymani-Mohammadi & Singer, 2013). The overall goal of this project is to demonstrate the use of BLI with the GS isolate, thereby eliminating the need for antibiotic treatment before and during mouse infection. We also demonstrated the stability of this GS strain as both circular and linear DNA are integrated into the genome. Finally, we compared measurements of the parasite burden of individual mice determined by bioluminescent imaging to parasite counts obtained from excised intestinal segments.

## RESULTS

### Modified *Giardia* GS strain enables the tracking of infection without the need for antibiotic treatment

To monitor infection dynamics *in vivo*, we infected C57BL/6J and BALB/c mice with GS-LucL and GS-LucC for 28 days. During the experiment, BLI was performed on days 0, 6, 8, 14, and 28 to measure infection burden and parasite clearance (Fig. 1). In the C57BL/6J mice, we recorded the highest bioluminescence levels for both GS-LucL and GS-LucC between days 6 and 8 post-infection (pi). Bioluminescence returned to baseline around day 10 and remained there through 28 dpi, indicating parasite clearance (Fig. 1A-D). Similarly, bioluminescence in BALB/c mice infected with either GS-LucL or GS-LucC was the highest at day 6, with a return to baseline at day 8 and continuing until euthanasia at day 28 (Fig. 1E-H). Although individual C57BL/6J mice exhibited varying infection kinetics, each animal reached a peak infection burden at 6 or 8 dpi (Fig. 1C-D). Peak infection at 6 dpi in the BALB/c mice was more consistent, with less variation among animals (Fig. 1G-H). The coat color of the mice influenced baseline bioluminescence: The C57BL/6J mice have black fur, and uninfected mice had slightly lower baseline radiance levels at day 0 compared with BALB/c mice that have white fur (Figure 1 C-D and G-H). Since radiance levels at the peak of infection were slightly higher in GS-LucC compared to GS-LucL, we utilized GS-LuC in the remaining experiments.

**Figure 1:**
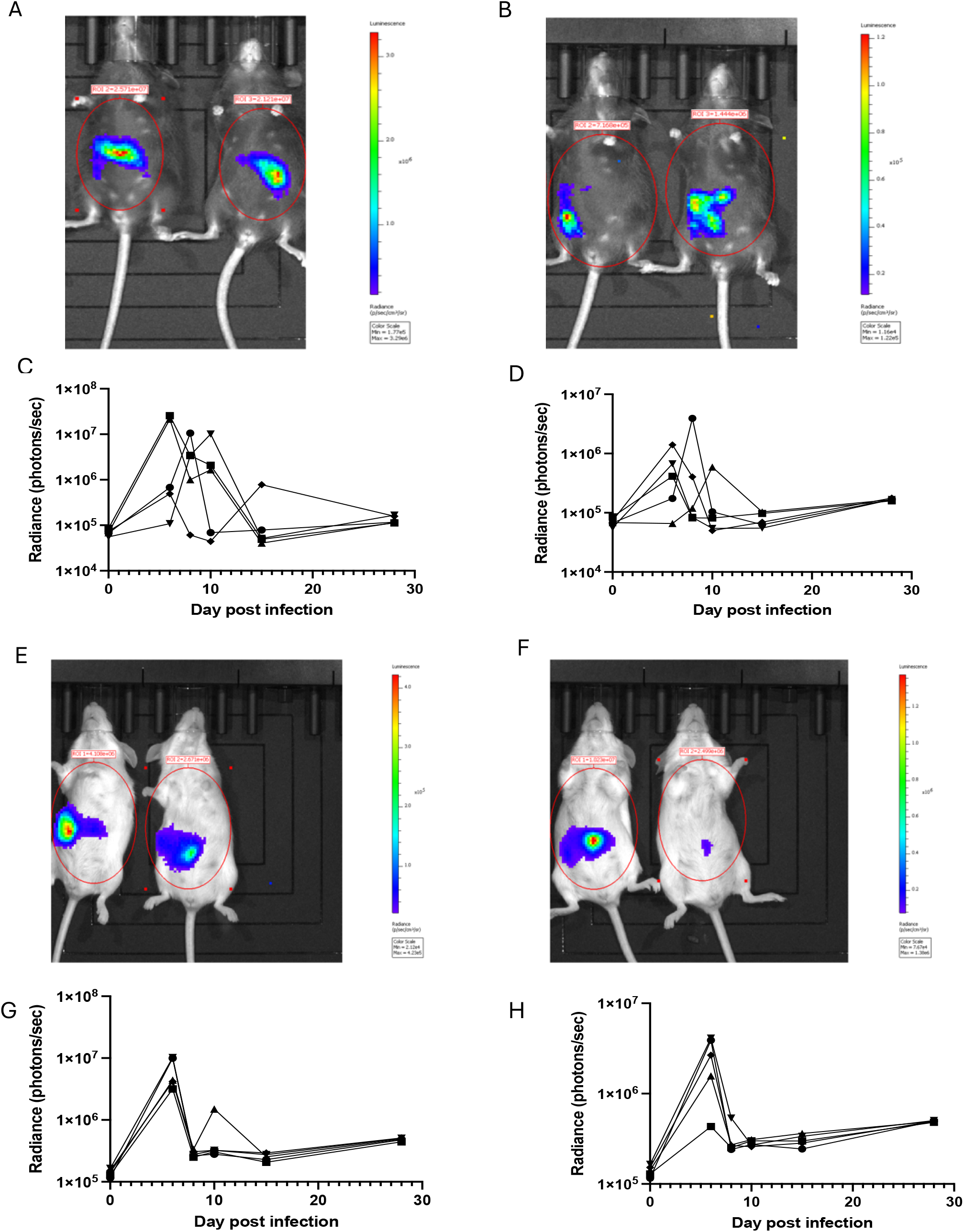
Infection dynamics of linear (GS-LucL) and circular (GS-LucC) *Giardia* in C57BL/6J and BALB /C wild-type mice. (A-D) Bioluminescent imaging of C57BL/6J mice infected with GS-LucC (A, B) and GS-LucL (C, D). Bioluminescent imaging of BALB/c mice infected with GS-LucC (E, F) and GS-LucL (G, H). Representative images are shown in panels A, B, E and F with radiance depicted on a variable color-scale adjusted to maximize the visual signals. Graphs in C, D, G and H indicate infection kinetics in individual mice imaged at multiple time-points throughout the 28 day experiment. Note that day 0 imaging was performed prior to mice being gavaged with trophozoites.

### Tracking stability of bioluminescence in immune-deficient mice

To verify that GS-Luc parasites remain stable in vivo without the loss of the FLuc transgenes, we infected immunodeficient BALB/c SCID mice with GS-LucC. These mice are unable to clear parasites (Singer et al., 1998). Infected animals were imaged at 6, 8, 13, and 21 dpi to track parasite burden. Unlike in wild-type BALB/c and C57BL/6J mice, the bioluminescent signal increased over the 21-day period (Figure 2). Comparing the average radiance between days 13 and 21 shows no difference between groups at these times. This confirms that the parasites are stable and retain their luciferase expression if the host immune system does not eliminate them. We also infected a separate group of immunodeficient C57BL/6J *Tcrb*^*tm1Mom*^ mice with previously constructed WB-Luc parasites (Barash, Nosala, et al., 2017). Like BALB/c SCID mice infected with GS-LucC, these mice exhibited high bioluminescence signals (between 10^8^ and 10^9^ photons/sec/cm^2^) for at least 21 days (Fig. 2F).

**Figure 2:**
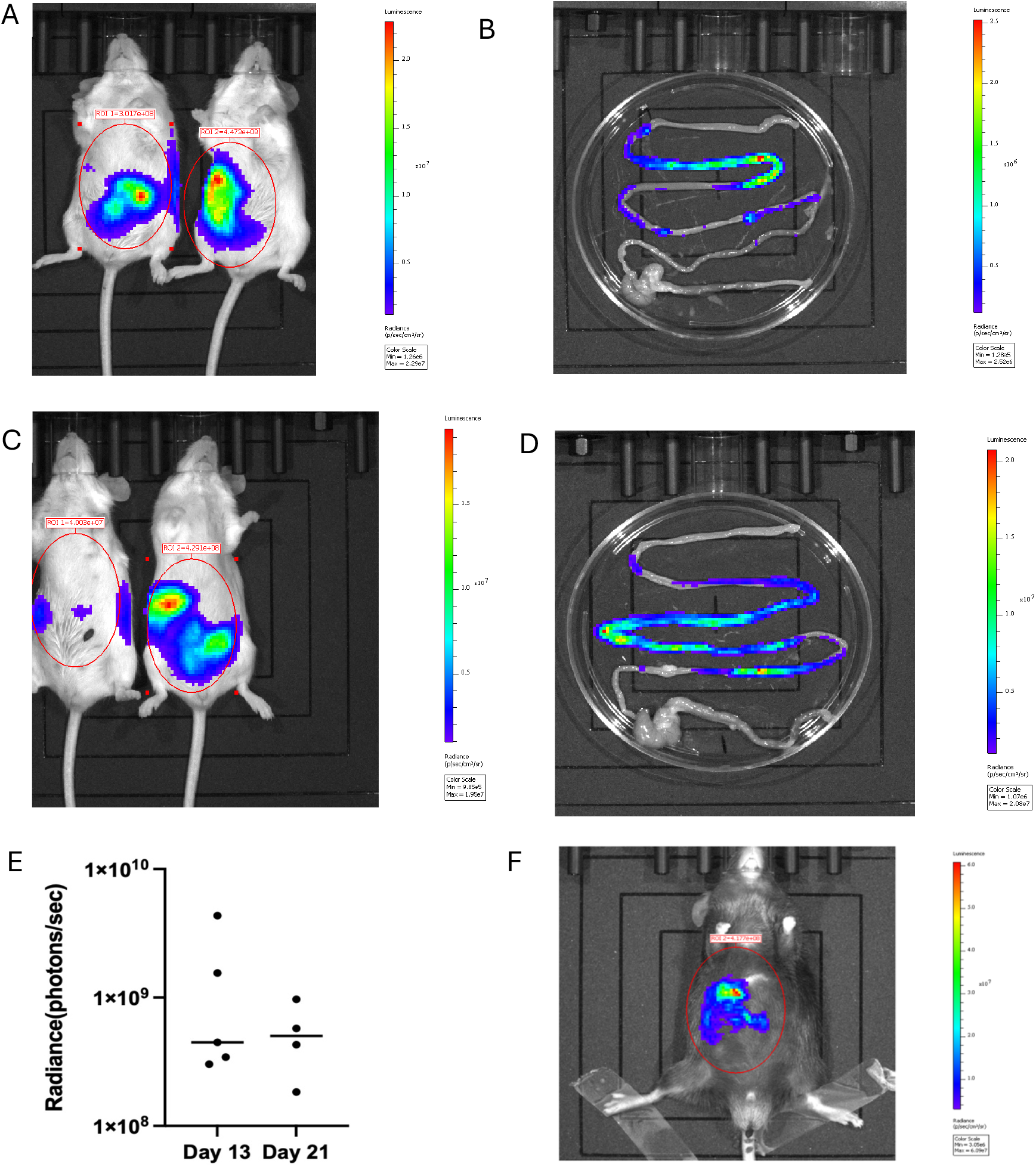
Persistence of bioluminescence in infections in immunodeficient mice by in vivo and ex vivo bioluminescent imaging. (A) Bioluminescent imaging at 13 dpi of a BALB/c SCID mouse infected with GS-LucC. (B) Excised intestinal tissue from the mouse imaged in (A). (C) Bioluminescent imaging at 21 dpi of a BALB/c SCID mouse infected with GS-LucC. (D) Excised intestinal tissue from the mouse imaged in (C). (E) Total radiance in individual BALB/c SCID mice on days 13 and day 21 post-infection with GS-LucC. No significant difference in radiance intensity was observed between days. (F) Bioluminescent imaging of a C57BL/6J *Tcrb*^*tm1Mom*^ mouse infected with the WB-Luc strain at 21 dpi.

### Bioluminescence intensity correlates with parasite counts

We next sought to determine whether bioluminescent imaging was an equally effective means to quantify infection burden compared with direct counting of intestinal trophozoites. We therefore infected mice with GS-LucC under conditions expected to produce a range of intestinal parasite burdens and counted trophozoite numbers in the small intestine immediately after performing bioluminescent imaging. We observed a strong positive correlation (r = 0.915, p < 0.001) between the observed radiance and the corresponding parasite counts from excised tissue sections (Figure 3). This supports the use of BLI with the GS-LucC strain as an alternative method to quantify parasite burden.

**Figure 3:**
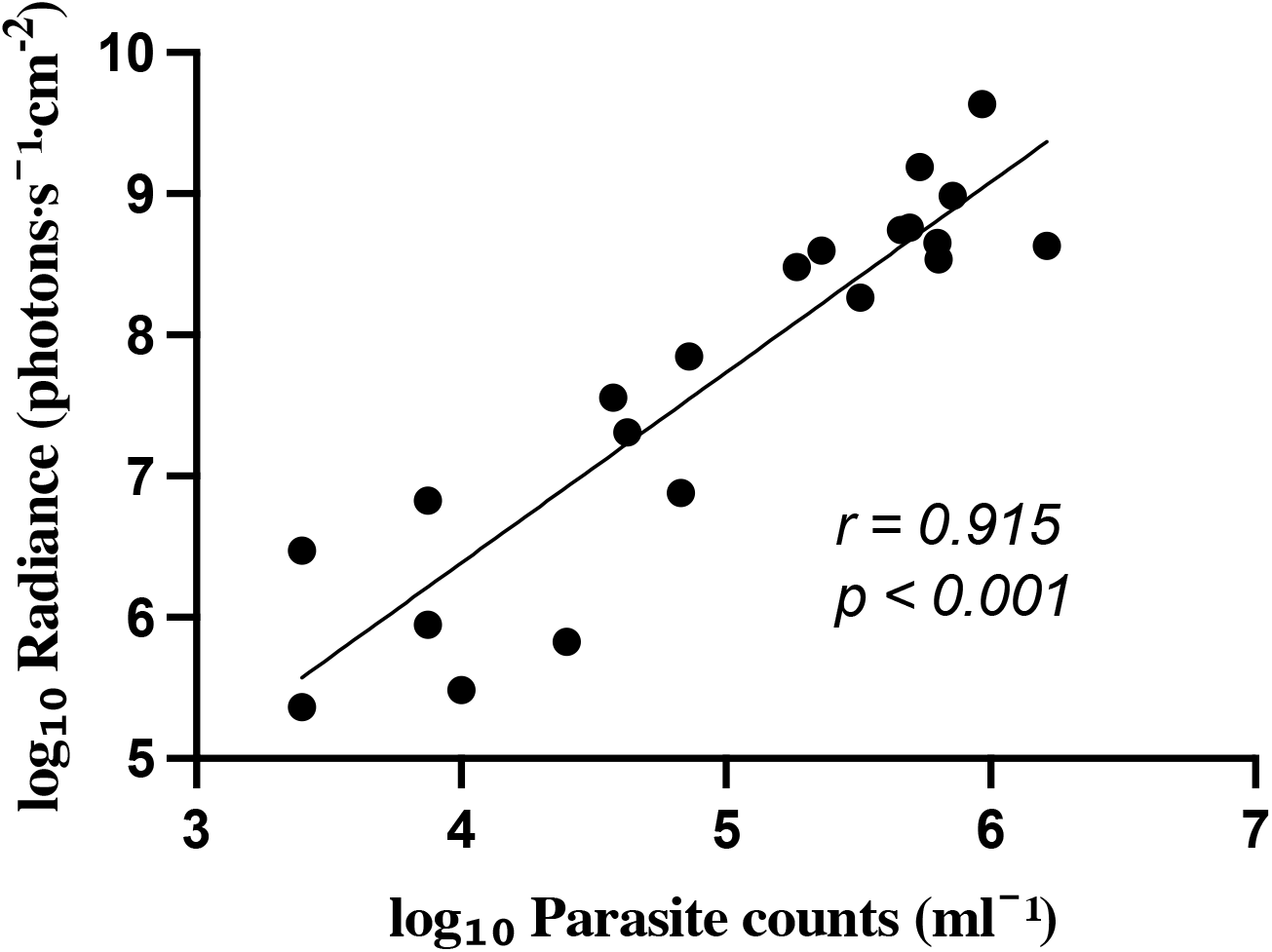
Scatterplot showing luminescence (radiance) as a function of parasite density. Mice were euthanized immediately after bioluminescent imaging and parasite numbers were determined in multiple (3-5/mouse) segments of the small intestine using a hemocytometer. The highest parasite burden from among all segments was assigned as the parasite burden for the scatter plot. Both variables were log_10_-transformed prior to plotting and calculation of a linear regression line. Data points represent different individual experiments using BALB/c or BALB/c SCID mice infected with GS-LucC (n = 20).

We also performed both BLI and manual counting on a number of mice where no parasites were observed by manual counting, suggesting there were fewer than 2,000 parasites/cm of small intestine. To determine a limit of detection for BLI, we calculated a threshold defined as the mean radiance of uninfected controls plus 2 standard deviations (mean + 2 SD), corresponding to 3.25 × 10^5^ photons/sec/cm^2^. The radiance values in the mice with uncountable parasites (n=9) remained just below this BLI threshold (range: 1.77 × 10^5^ to 2.97 × 10^5^ photons/sec/cm^2^), suggesting that whole-abdomen BLI has a very similar level of sensitivity as manual counting. These animals were excluded from the regression analysis because they had zero manual counts.

## DISCUSSION

The development of luciferase-expressing *Giardia* strains offers a robust, non-invasive method for tracking infection progression and host-parasite interactions in *Giardia-*infected mouse models within living organisms. In this study, we modified the GS strain to create two luciferase-expressing GS (GS-Luc) reporter parasites, thus enabling us to monitor infection patterns over time in both immunocompetent and immunodeficient mouse models. Our data revealed the colonization pattern of parasites in the mouse small intestine, as well as the stability and persistence of the parasites. This validated bioluminescent imaging (BLI) as a sensitive and quantitative approach for studying *Giardia* in mouse models, without the need for antibiotic treatment prior to infection or to euthanize animals at multiple timepoints to study infection kinetics.

In C57BL/6J and BALB/c mice, bioluminescence peaked between days 6 and 8 post-infection, followed by a rapid decline back to baseline around day 10. This pattern aligns with the known acute phase of infection and parasite clearance in *Giardia* murine models, during which immune responses, such as CD4+ T cell activation and IgA production, facilitate parasite clearance (Hill et al., 1983; Singer & Nash, 2000; Solaymani-Mohammadi & Singer, 2011). Bioluminescence remained at baseline up to day 28 post-infection, reflecting the self-limiting nature of *Giardia* infection. This infection kinetic is also in agreement with previous studies using manual trophozoite counts and stool PCR to quantify infection burden, further confirming the accuracy of BLI in murine infections (Barash, Nosala, et al., 2017; Bartelt et al., 2013).

Our findings build directly on the foundational work by Barash, Nosala, et al., (2017) who first established bioluminescent imaging as a powerful approach to visualize *Giardia* infection dynamics in vivo using the assemblage A strain, WBC6. Consistent with their observations, we show that bioluminescence provides a sensitive, longitudinal readout of parasite burden and correlates strongly with independent measures of infection intensity. However, our study extends this approach to the assemblage B GS isolate, which is more representative of human infections and, critically, does not require antibiotic pretreatment or gnotobiotic conditions to establish infection. While Barash, Nosala, et al., (2017) emphasized the spatial organization of parasites into high-density foci and the relationship between density and encystation, our work focuses on validating bioluminescence as a quantitative alternative to track infection burden in an immunocompetent host with an intact microbiome. The strong correlation between radiance and direct trophozoite counts supports the use of GS-Luc parasites as a robust, physiologically relevant tool for monitoring infection dynamics. The infection dynamics between the C57BL/6J and BALB/c mouse strains were similar, with slight variations among individual C57BL/6J mice. This could be due to immunological biases between mouse strains that may influence the timing of parasite colonization, as well as inflammatory responses (Bernal et al., 2021; Ferreira et al., 2018). Moving forward to other experiments, we used GS-LucC since it showed more stability in both mice strains as shown in Figure 1.

In immunodeficient C57BL/6J *Tcrb*^*tm1Mom*^ and BALB/c SCID mice, both luciferase-expressing *Giardia* strains, WB-Luc and GS-LucC, respectively, showed similar infection patterns. Starting on day 7 post-infection, we observed a steady increase in radiance without decline throughout the infection period. This confirms previous findings that these animals are unable to clear parasites or control infections (Byrd et al., 1994; Singer & Nash, 2000). The prolonged bioluminescent signal suggests these modified *Giardia* strains remain stable and can effectively colonize and replicate in the mouse intestine. Additionally, luciferase expression continues without continuous antibiotic selection, making it suitable for chronic infection studies in mice. Notably, GS-LucC showed no differences in infectivity, luminescence, or stability. The positive correlation between radiance and parasite counts confirms that photon emission from GS-Luc parasites serves as a reliable way for quantifying in vivo infection burden. This validates BLI as a non-invasive method for monitoring the progress and clearance of *Giardia* infections. The correlation between radiance and parasite numbers shows that luciferase expression remains stable during infection and radiance increases proportionally with parasite density, consistent with a previous study (Barash, Nosala, et al., 2017). While stool PCR and manual trophozoite counts have been widely used to estimate *Giardia* burden, these approaches are limited by intermittent stool shedding and uneven parasite distribution along the small intestine, which can introduce sampling error (Barash, Nosala, et al., 2017; Duque-Beltrán et al., 2002; Fink et al., 2020). In contrast, BLI enables longitudinal, whole-abdomen assessment of infection burden within the same animal, reducing variability associated with terminal or region-restricted measurements.

Minor variations observed in these experiments may be due to BLI detection limits or biological factors, such as substrate circulation and oxygen levels, which can independently affect luciferase activity (Andreu et al., 2011). Also, BLI is generally limited by abdominal photon attenuation, as the coiling of the small intestine increases scattering and overlying organs such as the liver absorb emitted light, reducing detection from deeper intestinal regions (Contag et al., 1997; Jacques, 2013).

These findings highlight the dual utility of the GS-Luc parasites, as they enable real-time visualization of *Giardia* infection dynamics without significantly altering the intestinal microbiota caused by antibiotic treatment prior to mice infection (Barash Maloney, et al. 2017). It is also a reliable measure for tracking parasite burden longitudinally within the same animal, thereby reducing experimental variability and the need for multiple animals. Overall, BLI using GS-Luc in mice appears more sensitive than ex vivo direct counts, as it captures a larger surface area than counting from a portion of the small intestine.

Future research should utilize this imaging system to explore the cellular and molecular processes involved in parasite clearance and persistence. Moreover, applying this method to investigate coinfections, nutrient effects, or microbiota composition could offer a new understanding of the complex factors in *Giardia* pathogenesis. Beyond in vivo studies, GS-Luc parasites could be suited for in vitro drug screening and real-time quantification of parasite growth in co-culture systems with intestinal epithelial cell lines or organoids, enabling mechanistic studies under controlled conditions.

## MATERIALS AND METHODS

### *Giardia* strains, growth media

*Giardia duodenalis*, GS strain was obtained from the ATCC (#50581). WB-luc (WBC6/C2) that express the *P. pyralis* (firefly) luciferase gene (Barash et al., 2017) were obtained from Scott Dawson (University of California, Davis). Parasites were cultured in modified TYI-S-33 medium supplemented with bovine bile (Sigma-Aldrich), 5% fetal bovine serum (Invitrogen), 5% adult bovine serum (Invitrogen) and antibiotic plus antimycotic (Invitrogen) in sterile 16.5 mL screw-capped tubes (Fisher Scientific) and incubated on a slant at 37°C.

### Plasmid construction Cloning of TPI into pUC19

To create pUC-TPI, theTPI1 locus was amplified from *Giardia* GS genomic DNA with Q5® Hot Start High-Fidelity DNA Polymerase (NEB) using 5’-AGA GGA TCC CTT CTA CAC ATT GAT CAG CGC CC-3’ (forward) and 5’-CG GTA CCC CCA TTG AGG AGA CCC AGA G-3’ (reverse) as primers. The pUC 19 (New England Biolabs) plasmid backbone was amplified using the primers 5’-T CAA TGG GGG TAC CGA GCT CGA ATT CAC-3’ (forward) and 5’-GT GTA GAA GGG ATC CTC TAG AGT CGA CCT-3’ (reverse). TPI PCR products were treated with DpnI restriction endonuclease (New England Biolabs) and were purified with Monarch PCR and DNA Cleanup Kit (New England Biolabs) in parallel with pUC19 PCR products. PCR products were ligated using the Gibson Assembly as described previously (Gibson et al, 2009) and transformed into *E. coli* DH5α (NEB). The resulting plasmid was verified by whole plasmid sequencing (Plasmidsaurus; Eugene, OR).

### Insertion of FLuc/pac cassette into TPI locus (Cloning of CWPLuc into pUC-TPI)

CWP1-Luc DNAwas a gift from Dr. Scott Dawson, UCDavis (Barash et al., 2017). The pUC-Tpi plasmid was amplified with Q5® Hot Start High-Fidelity DNA Polymerase (NEB) using the primers 5’-CTG CTG TAA GAC GTC ATA CGG TAT GGC TAA CTC A-3’ (forward) and 5’-GT TAG CCA GCT CCT TTG TTA TTC TTC AGT GTA TGG ATT GAC A-3’ (reverse). In parallel, the portions of CWP1-Luc encoding firefly luciferase and the puromycin resistance cassette were amplified using the primers 5’-CAA AGG AGC TGG CTA ACA GTC TAC AGT CTA CAA TTT ACA GTA-3’ (forward) and 5’-G TAT GAC GTC TTA CAG CAG TTC GGG AAG TTT CCT-3’ (reverse). PCR products were cloned as described above for pUC19-TPI. The resulting plasmid was named ***pTP-CWPLuc*** (supplemental Figure 1).

### Transfection and selection

#### pTP-CWPLuc

was linearized using a BamHI restriction digest. Both linear and circular plasmids were transfected independently into GS by electroporation (~20 ug and 40 ug DNA respectively) as previously described (Singer et al., 1998). Strains were initially selected with 10 ug/ml puromycin for ~7 days and then expanded with 50 ug/ml puromycin. After cells were passaged once and frozen stocks prepared, puromycin was no longer included in the media. The parasite line derived from transfection with circular plasmid was denoted GS-LucC and the line derived with linearized DNA was denoted GS-LucL.

### Animal infections

Six-to eight-week-old BALB/c, C57BL/6J, C57BL/6J *Tcrb*^*tm1Mom*^ and CBySmn.Cg-*Prkdc*^*scid*^/J (BALB/c SCID) mice were purchased from Jackson laboratory (Bar Harbor, ME). Animals were housed and maintained in specific-pathogen-free conditions. Animals were infected and imaged at intervals up to 28 days. Trophozoites were detached on ice and washed with cold PBS before counting using a hemocytometer. Each mouse was infected by oral gavage of 1 × 10^6^ trophozoites in 100 μL of sterile PBS (Fink et al., 2020). All animal experiments were performed according to IACUC-approved protocols.

### Live imaging

To image *Giardia* colonization of the mouse gut, we used the luciferase substrate, IVISbrite D-Luciferin Potassium Salt Bioluminescent Substrate (10 x 1g) (XenoLight) from Revvity at 150 mg/kg body weight. The substrate was intraperitoneally injected, and the mouse was left to rest for 15 minutes to allow the substrate to circulate. Each mouse was then placed in an isoflurane (2 – 3%) chamber to be sedated before imaging using an IVIS chamber (Caliper life sciences).

### Manual parasite counts

Immediately after imaging, mice were euthanized, and 5 cm sections were taken from the duodenum, jejunum, and ileum and placed in cold PBS. The sections were opened longitudinally before being minced in 5 mL of cold PBS. Parasite numbers were counted using a hemocytometer, resulting in a detection limit of 2,000 trophozoites/cm.

## ACKNOWLEDGMENTS

This work was supported by National Institutes of Health (NIH) grants R15AI109591 and R21AI166467 to Steven M Singer. Rita T Kosile was supported by NIH 5F31AI183732-02, and the Georgetown University Preclinical Imaging Research Laboratory (PIRL) was supported by NIH/NCI CCSG grant P30-CA051008. The funders had no role in study design, data collection, and interpretation, or the decision to submit the work for publication.

---

**Supplemental Figure 1.**
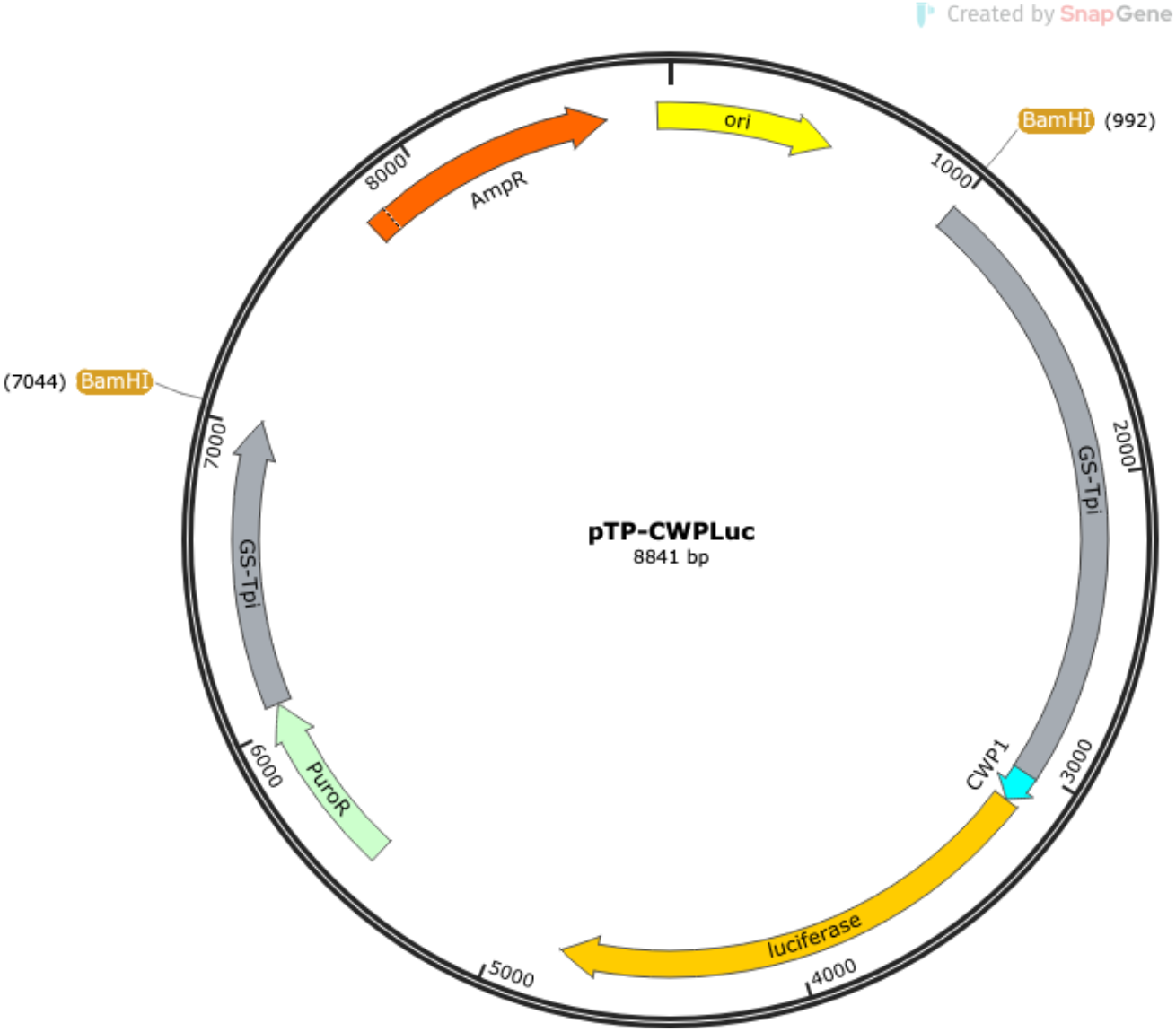
Map of pTP-CWPLuc used to integrate the firefly luciferase gene into the TPI locus of *Giardia* isolate GS.

